# Tumor Edge-to-Core Transition Promotes Malignancy in Primary-to-Recurrent Glioblastoma Progression in a PLAGL1/CD109-mediated mechanism

**DOI:** 10.1101/2020.09.14.293753

**Authors:** Chaoxi Li, Hee Jin Cho, Daisuke Yamashita, Moaaz Abdelrashid, Qin Chen, Soniya Bastola, Gustavo Chagoya, Galal A. Elsayed, Svetlana Komarova, Saya Ozaki, Yoshihiro Ohtsuka, Takeharu Kunieda, Harley I Kornblum, Toru Kondo, Do-Hyun Nam, Ichiro Nakano

## Abstract

**Background:** Glioblastoma remains highly lethal due to its inevitable recurrence. This recurrence is found locally in most cases, indicating that post-surgical tumor-initiating cells (TICs) accumulate at tumor edge. These edge TICs then generate recurrent tumors harboring new core lesions. Here, we investigated the clinical significance of the edge-to-core transition (ECT) signature causing glioblastoma recurrence and sought to identify central mediators for ECT.

**Methods:** First, we examined the association of the ETC-related expression changes and patient outcome in matched primary and recurrent samples (n=37). Specifically, we tested whether the combined decrease of the edge TIC marker PROM1 (CD133) with the increase of the core TIC marker CD109 representing ECT during the primary-to-recurrence progression indicates poorer patient outcome. We then investigated the specific molecular mediators that trigger tumor recurrence driven by the ECT signature. Subsequently, the functional and translational significance of the identified molecule was validated within our patient-derived tumor edge-TIC models *in vitro* and *in vivo*.

**Results:** Patients exhibiting a CD133^down^/CD109^up^ signature during recurrence representing ECT displayed a strong association with poorer progression-free survival and overall survival among all tested patients. Differential gene expression identified that PLAGL1 was tightly correlated with the core TIC marker CD109 and was linked to a shorter survival of glioblastoma patients. Experimentally, forced PLAGL1 overexpression enhanced, while its knockdown reduced, the glioblastoma edge-derived tumor growth *in vivo* and subsequent mouse survival, suggesting its essential role in the ECT-mediated glioblastoma development.

**Conclusions:** ECT is likely an ongoing lethal process in primary glioblastoma contributing to its recurrence partly in a PLAGL1/CD109-mediated mechanism.

**Key Points:**

1. ECT is a pathobiological process contributing to glioblastoma lethality
2. The CD133^down^/CD109^up^ signature is a novel prognostic molecular biomarker in ECT
3. PLAGL1 regulates growth of edge-located tumor-initiating cells

**Importance of the Study:** Very few studies have sought to longitudinally characterize the transition of molecular landscapes from primary to recurrent glioblastoma. Post-surgical edge-located TICs are presumably the predominant source of tumor recurrence, yet this cellular subpopulation in glioblastoma remains largely uncharacterized. This study evaluates the significance of glioblastoma edge-derived core transition (ECT) for tumor recurrence in the primary-recurrent paired sample set. We elucidate a prognostically-significant shift in molecular and cellular phenotypes associated with ECT in the CD133^down^/CD109^up^ group. Moreover, our results provide clinical and experimental evidence that PLAGL1 is a mediator of glioblastoma ECT and its subsequent tumor development by the direct transcriptional regulation of the core TIC marker CD109.

## Introduction

Glioblastoma is an incurable universally lethal disease^1^ and characterized by inter- and intra-tumoral heterogeneity^2–6^. Transcriptome-based subtyping of individual tumors is considered a milestone discovery of the past decade^7,8^; nonetheless, this molecular subtyping has yet to change clinical management, unlike other cancers that now have distinct treatment options instructed by particular genetic subtype information (e.g. breast cancer, neuroblastoma)^9,10^. In sharp contrast to the accumulating experimental evidence for the mesenchymal shift of glioblastoma tumors being tightly associated with a gain of malignancy and therapy resistance in various model systems, clinical data remains lacking to suggest that mesenchymal glioblastoma gains benefit from more extensive and/or different therapies. In addition, multiple independent large-scale studies have clarified that the transcriptomic subtype switch between primary and recurrent glioblastomas is simply a random event without any clear trend of one way or the other including toward the mesenchymal shift^11^.

Most glioblastomas recur within a few years as the main cause of its dismal prognosis in the developed countries^12^. The large degrees of molecular difference between primary and recurrent tumors have been recognized by various OMICs analyses including deep sequencing, both with tumor tissues^13,14^ and at the single cell level^15^. Since the brain tissues adjacent to surgical resection are the most frequent sites of tumor recurrence, the normal parenchyma-tumor core interface (tumor edge) presumably contains post-surgical tumor-initiating cells (TICs; also termed recurrence-initiating cells) after craniotomy. Molecular and, more importantly, phenotypic characterization of these edge-TICs may lead to the identification of a means to inhibit the process of tumor recurrence following craniotomy.

Diffuse infiltrative glioblastomas, when they recur, are detected by the propagation of new tumor core lesions, indicating the edge-to-core transition (ECT) is likely critical step toward patient lethality. Nonetheless, these lethal seeds for tumor recurrence are mostly, if not entirely, surgically-untouchable due to the presence of intermingled normal functional brain cells including neurons. In fact, despite recent advances in surgical technology increasing the extent of resection of the core lesion with the neuro-radiological confirmation of nearly 100% resection of the enhancing abnormality on post-operative MRI, the improvement of post-surgical patient survival remains marginal. Therefore, further attention needs to be placed on the remaining edge lesions (T2/FLAIR abnormality without Gadolinium enhancement on MRI) and ECT during recurrent tumor development as a clinically-significant consequence of treatment failure to glioblastoma. In order to uncover the functional roles of tumor cells within this edge microenvironment, our recent studies have undertaken a program to isolate and characterize regionally-distinct tumor cell populations by using awake surgery to obtain reasonable amounts of edge tissues without harming patients, allowing us to functional identify CD133 and CD109 as the representative molecules to mark the *edge*-located and *acquired core*-associated TICs, respectively^3,16–18^.

In the current study, we investigated this presumptive transition of CD133^high^/CD109^low^ cells to CD133^low^/CD109^high^ cells as the representative of highly-lethal ECT dynamics by using 37 pairs of samples from matched primary and recurrent glioblastoma tumors. We then postulated that the decline of CD133 expressing TICs and the increase of CD109-expressing TICs indicates active ECT progression, worsening the patients’ prognosis. To test this idea, we segregated our longitudinal sample set into two groups based on the CD133/109 expression changes. A set of integrated multimodal analyses was performed, followed by the pre-clinical validation of the identified molecular target as a functional key determinant for ECT-related glioblastoma aggressiveness.

## Materials and Methods

### Patients, Specimens, and Ethics

All 37 longitudinal glioblastoma cases were treated at Samsung Medical Center and Seoul National University Hospital and the tumor tissues were collected for research under the approved institutional review boards. Detailed methods are described in the previous study^19^ and Supplementary material. For the pre-clinical studies, four patient-derived glioma sphere models were used, including three pair of tumor core- and edge-derived ones (1051E and C, 1053E and C, 0573E and C) as well as one tumor edge-derived sphere line (101027E), which were established and described elsewhere^3,16–18,20–22^. In short, with the signed patient consent, the senior author (IN) performed supra-total resection of glioblastoma tumors under the awake setting and resected both tumor core (T1-Gadolinium(+) tumors) and edge (T1-Gadolinium(−)/T2-FLAIR abnormal tumors in the non-eloquent deep white matter) to achieve maximal tumor cell eradication without causing any permanent major deficit in the patients (Supplementary Fig.1A). After the confirmation of enough tumor tissues from both lesions secured for the clinical diagnosis, remaining tissues were provided to the corresponding scientists following de-identification of the patient information. Both the core-derived and edge-derived glioma spheres were established in the same culture condition ^3,16–18,20–23^. and their spatial identities, termed *core-ness* and *edge-ness,* were confirmed by a set of xenografting experiments into mouse brains (details described in ^18^). Only those that passed this confirmation were used for this study. The other patient-derived glioma sphere models were established as “core-like glioma spheres” using the same protocol and reported elsewhere^18^. All these patient-derived glioma models were periodically checked with the mycoplasma test and the Short Tandem Repeat (STR) analysis. All work related to pre-clinical data was performed under an Institutional Review Board (IRB)-approved protocol (N150219008) compliant with guidelines set forth by National Institutes of Health (NIH).

### Public Microarray Data Processing

Three RNA sequencing datasets were downloaded from the Gene Expression Omnibus database(https://www.ncbi.nlm.nih.gov/geo/), including GSE63035, GSE67089 and GSE113149^17,23,24^. RNA sequencing data of 29 longitudinal samples are derived from GSE63035, and 8 longitudinal samples are newly added, all the samples are IDH-wild type. The GSE67089 datasets contained gene expression data of MES, PN glioma sphere cells and Neuron progenitor cells. The GSE113149 included the microarray data for sh-NT versus sh-CD109 in glioblastoma sphere 267. The RNA sequencing data of TCGA database was acquired from the TCGA Research Network (https://www.cancer.gov/tc-ga.) and visualized by Gliovis^25^ (http://gliovis.bioinfo.cnio.es/).

### *In vitro* experiments

Detailed methods are described in the Supplementary material.

### *in vivo* mouse experiments

All animal experiments were performed at UAB under the Institutional Animal Care and Use Committee (IACUC)-approved protocol according to NIH guidelines. Detailed methods are described in the Supplementary material.

### Statistical Analysis

All data are presented as mean ± SD. The number of replicates for each experiment was stated in Figure legends. Statistical differences between two groups were evaluated by two tailed *t*-test. The statistical significance of Kaplan–Meier survival plot was determined by log-rank analysis. A statistical correlation was performed to calculate the regression R^2^ value and Pearson's correlation coefficient. Statistical analysis was performed by Prism 8 (GraphPad Software), unless mentioned otherwise in figure legend. P < 0.05 was considered as statistically significant.

## Results

### Patients in CD133^down^/CD^109^up group exhibit worse prognoses with a trend towards an increased mesenchymal signature

Based on our previous data^17,18^, we used CD133 mRNA and CD109 mRNA to indicate edge-ness and core-ness, respectively, a concept that we validated with 19 paired GBM edge- and core-samples**(Supplementary Fig. 2)**. We reasoned that the loss of CD133 mRNA (CD133^down^) and gain of CD109 (CD109^up^) were indicative of the edge-to-core transition in glioblastoma. Based on the differential RNA expression profiles as determined by RNA-sequencing (seq) of 37 primary and recurrent glioblastoma pairs, 15 patients were assigned to the CD133^down^/CD109^up^ group as representative of ECT, while the other 22 patients were assigned as control arms (Others,either CD133^down^/CD109^down^, CD133^up^/CD109^down^, or CD133^up^/CD109^down^) for comparison. Both groups displayed similar average age, sex, distant recurrence profiles, and post-surgical therapy regimens. **(Table 1, Supplementary Table 1).**We then investigated the progression-free survival and overall survival in these four groups. The CD133^down^/CD109^up^ group exhibited a substantially worse progression-free survival (*p*=0.024) and overall survival *(p*=0.043) compared with others**(Fig. 1A)**. Consistent with recent studies, both primary and recurrent tumors showed no significant difference in proportion among the three transcriptomic subtypes^6^. However, there was a trend that CD133^down^/CD109^up^ group was enriched in tumors of the mesenchymal subtypes upon recurrence (*p*=0.028) **(Fig. 1B)**. Nonetheless, in this patient cohort, the mesenchymal-ness of either primary or recurrent tumors did not show statistically-significant differences in prognosis. These findings suggested a significant association between the CD133^down^/CD109^up^ signature representing ECT and poorer patient prognoses, associated with increase of the mesenchymal subtype in the primary-to-recurrent glioblastoma progression.

**Figure 1.**
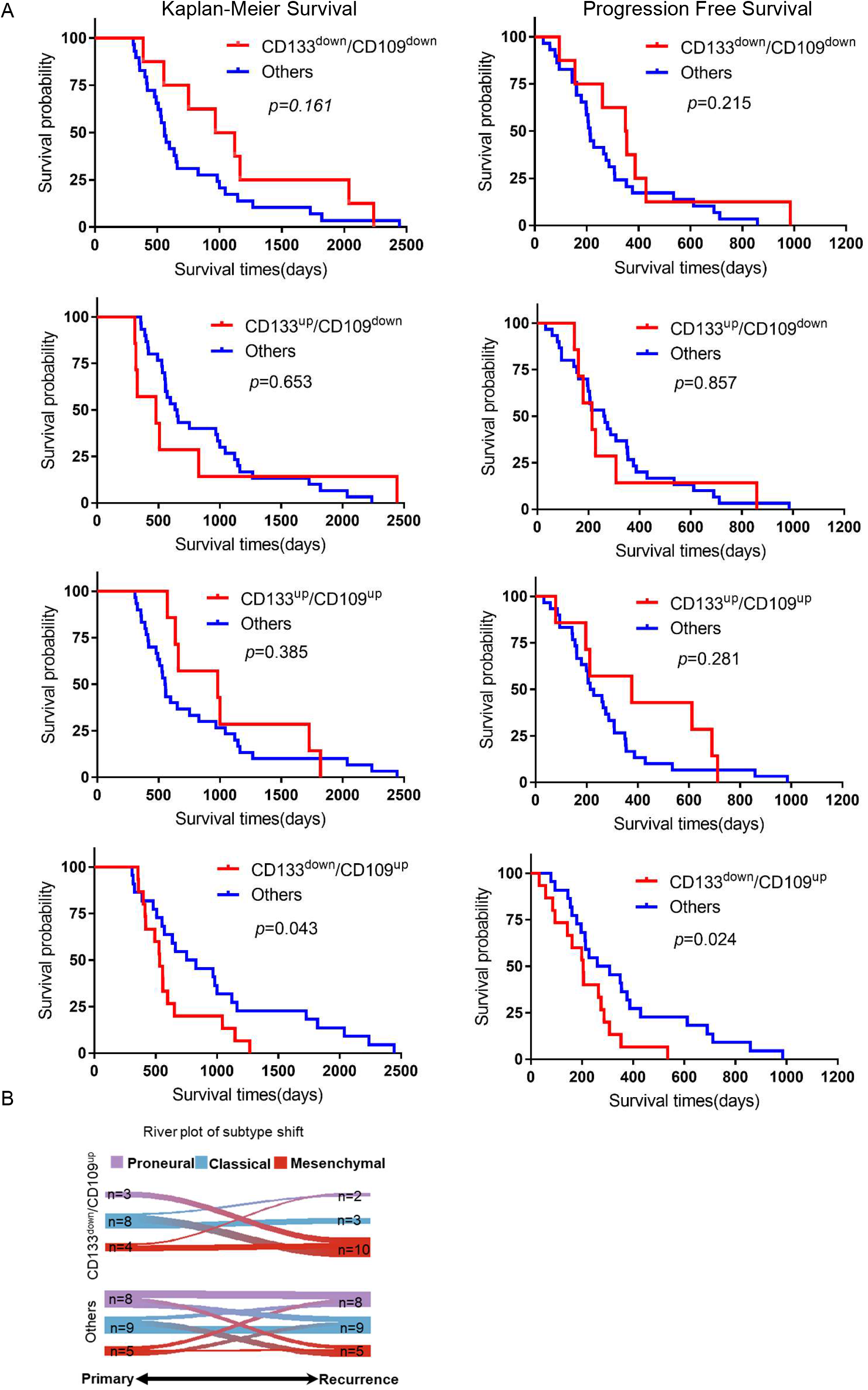
CD133^down^/CD109^up^ group exhibits worse prognoses accompanied by an increase in mesenchymal signature. (A) Kaplan-Meier analysis of Overall Survival (left) and Progression-Free Survival (right) of glioblastoma patients in CD133^down^/CD109^down^, CD133^up^/CD109^down^, CD133^up^/CD109^up^, and CD133^down^/CD109^up^ (red, top to bottom) with each collected remaining cases (Others, blue). (Log-rank test). (B) River-plot analysis of the molecular subtype shifts from primary to recurrence in CD133^down^/CD109^up^ (upper) and others (lower). (*p* =0.028, Chi square test)

### Longitudinal RNA-seq analysis identifies the differential expression profile associated with ECT including PLAGL1 and CD109

Next, we pursued a stepwise approach to identify a molecular target or targets that could mediate the observed molecular and phenotypic dynamics of ECT. First, we established a data analysis pipeline using all expressed genes in the RNA-seq data of the 37 longitudinal cases (n=22,255) **(Fig. 2A)**. Differential gene expression analysis identified 26 genes distinctively associated with the CD133^down^/CD109^up^ changes **(Supplementary Table 2)**. Unsupervised hierarchical clustering of those genes (n=155) segregated our cohort sample (n=37) into two distinctive subgroups (up- and down-regulated) **(Fig. 2B, C)**. In order to further elucidate the essential molecules governing ECT, we designed an integrated second step approach to evaluate the expression of these 26 up-regulated genes in our well-characterized glioma sphere models treated with either shRNA-based gene silencing of CD109 or flow cytometry to isolate CD109(+) cells. To this end, we used our recently-published RNA-seq data with two well-characterized tumor core-like glioma sphere models; g267 for shRNA and g1005 for flow cytometry^17^. As a result, *PLAGL1* was identified as being the gene whose expression most strongly correlated with that of *CD109* (FC>1.5, *p*<0.05) **(Fig. 2A, D, Supplementary Fig. 3A, B)**. Consistently, Pearson’s correlation analysis of the 37 glioblastoma paired samples indicated a strong linear relationship between *CD109* and *PLAGL1* relative expression (r = 0.7, *p*< 0.05) **(Fig. 2E)**. This *CD109-PLAGL1* expression correlation was also observed in four clinical datasets (TCGA, Rembrandt, CGGA, and CGGA GBM datasets) **(Fig. 2F).** qRT-PCR with two additional edge- and core-derived glioma sphere models (Edge- and Core-derived g1053 spheres and g0573 spheres) showed that both PLAGL1 and CD109 were higher in the core-derived, yet CD133 was up in the edge-derived, glioma spheres *in vitro* **(Fig. 2G).**

**Figure 2.**
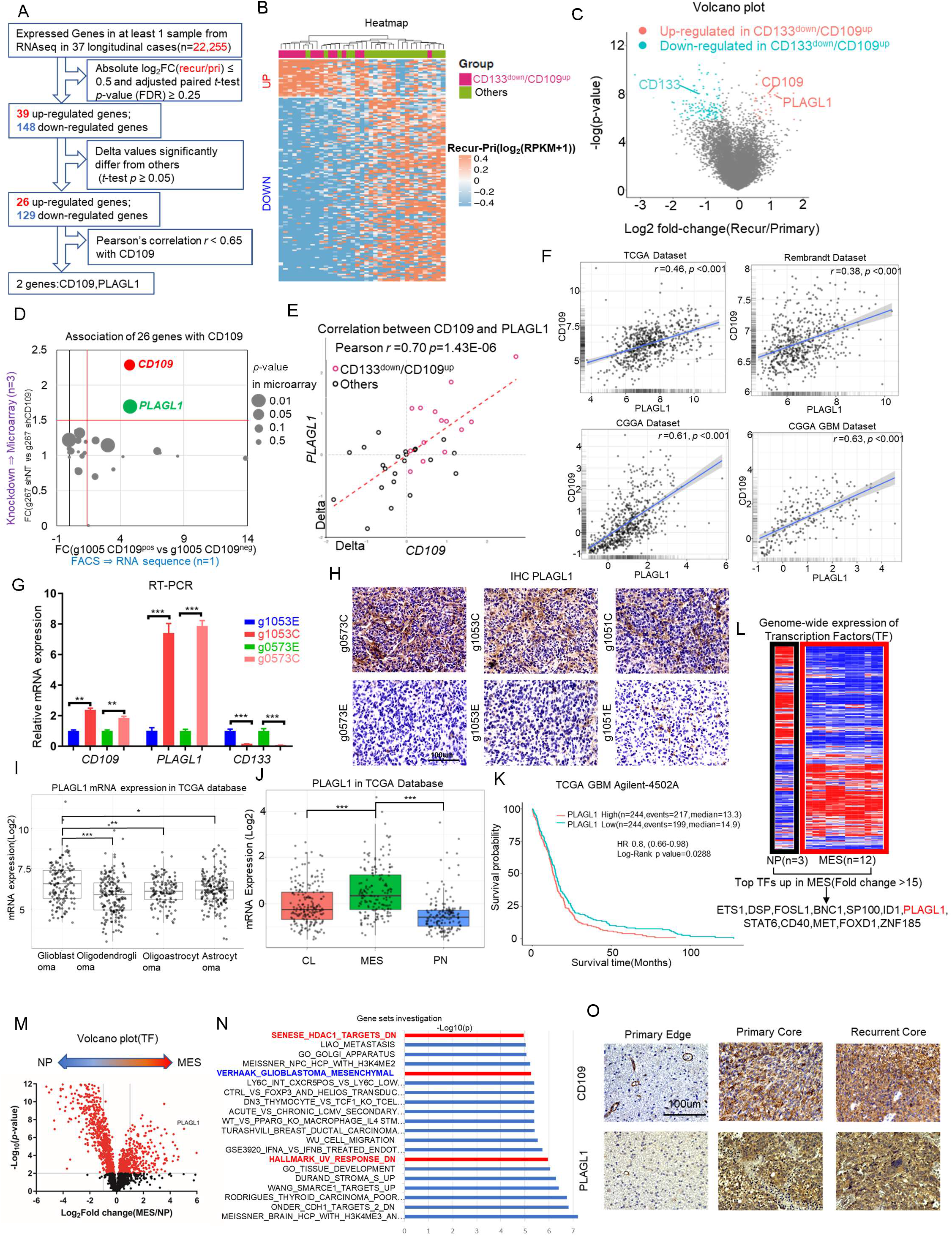
Longitudinal RNA-seq analysis identifies the differential expression profile associated with ECT including PLAGL1 and CD109. (A) Schematic demonstration of the filtering procedure of *PLAGL1* from 22,255 genes. (B) Heatmap depicting supervised hierarchical clusters of up- and down-regulated genes within recurrent glioblastomas of CD133^down^/CD109^up^ and others. (C) Volcano plot of RNA-seq data comparing CD133^down^/CD109^up^ and others. Red and blue dots refer to up- and down-regulated genes in CD133^down^/CD109^up^ group respectively. (D) Scatterplot comparing expression profiles of the 26 genes from RNA-seq analysis results within our glioma sphere models; MES-g1005 (n=1) FACS-sorted into CD109 negative to positive cells and represented along the x-axis (left to right respectively), while microarray relative expression of *CD109* in MES-g267 (n=3) with shNT and shCD109 is represented along the y-axis (down- and up-ward, respectively). (E) Scatterplot displaying the linear correlation between *CD109* and *PLAGL1* expressions in the 37 longitudinal cases. Pearson correlation coefficient (r) = 0.70 and *p*= 1.43E-06. (F) Scatterplot displaying the linear correlation between *CD109* and *PLAGL1* expressions in 4 public databases (TCGA, Rembrandt, CGGA, CGGA GBM), based on Pearson correlation test. (G) Bar graph displaying qRT-PCR results for the expression of *CD109, PLAGL1*, and *CD133* within edge- and core-derived sphere culture models of 2 glioblastoma patients (g1053 and g0573). Data are means ± SD (n=3). \*\*\**p*<0.001. (H) Representative images of immunohistochemistry (IHC) for PLAGL1 in mouse orthotopic xenografts with tumor core- (C) and edge(E)-derived glioma sphere models from 3 patients (g0573, g1053, and g1051). Scale bar 100um. (I) Boxplot diagram demonstrating PLAGL1 relative mRNA expression profiles from TCGA database across different gliomas subtypes. *p<0.05, **p<0.01, and *** *p*<0.001. (J) Boxplot diagram comparing relative expression profiles of *PLAGL1* among the 3 molecular subtypes (CL, MES, and PN) of glioblastoma within TCGA database. \*\*\**p*<0.001. (K) Kaplan-Meier survival curve of glioblastoma patients in the TCGA database. Patients were categorized into a ‘‘high’’ or ‘‘low’’ expression group based on the median *PLAGL1* expression in the Agilent 4502 microarray. (L) Heatmap of displaying expression profiles of transcription factors (TF) (n=2,766) across 4 MES glioma sphere lines compared with the neural progenitor sphere line (NP) (n=3 for each cell line). (M) Volcano plot comparing TF gene expressions (n=2,766) across MES and NP lines, highlighting *PLAGL1* (N) Up-regulated pathways in recurrent glioblastomas in CD133^down^/CD109^up^ group. (O) Representative IHC images for CD109 and PLAGL1 in primary edge and core, and recurrent core tumor tissues. Scale bar 100um.

To prospectively assess PLAGL1 localization in experimental tumors, we injected edge- or core-derived glioma spheres from 3 patients into immunodeficient mice. PLAGL1 showed its preferential expression in the tumor core-derived lesions (patient n=3) **(Fig. 2H)**. In the patient tumor data in TCGA, *PLAGL1* mRNA expression was relatively higher in glioblastomas compared to lower grade gliomas **(Fig. 2I).**In glioblastoma, *PLAGL1* mRNA was enriched in mesenchymal tumors **(Fig. 2J)**. As expected, glioblastoma patients with higher PLAGL1 expression exhibited shorter survival in the TCGA database **(Fig. 2K)**.

Since the *PLAGL1* gene encodes for C2H2 zinc finger (ZF) transcription factors (TFs)^26^, we sought to further confirm our results by cross-referencing them to our previously-established cDNA microarray dataset with the sphere lines established from either human neonatal brains (neural progenitors: NPs) or glioma patients with mesenchymal or core-like signature^23^. Among 2,766 human TFs^27^, 12 TFs, including *PLAGL1*, were highly overexpressed (fold change >15) in mesenchymal or core-like glioma sphere lines as opposed to NP counterparts (*p*<0.001) **(Fig. 2L)**, moreover, *PLAGL1* was the second highest C2H2-ZF TFs in MES cells **(Supplementary Fig.4)**. Consistently, a volcano plot displayed *PLAGL1* as a significantly upregulated gene in mesenchymal or core-like glioma spheres **(Fig. 4M).** Gene set enrichment analysis (GSEA) using the 26 upregulated genes identified their association with “HDAC1 targets” and “UV response DNA damage”, both of which our recent studies have identified as pathways tightly correlated to CD109-driven signals in glioblastoma and their TIC models **(Fig.4N)^17,18^**. Finally, we explored the expression of PLAGL1 in primary GBM edge, core lesions as well as their subsequent recurrent core tissues, which showed PLAGL1 higher in the core lesions in both primary recurrent tumors **(Fig. 4O)**. Collectively, these clinical and experimental data suggested PLAGL1 is possibly one key regulator in the edge-TICs to cause tumor core development in glioblastoma.

### Genetic perturbation of PLAGL1 reveals its role in glioblastoma tumorgenicity in the edge-TIC models

Following the identification of PLAGL1 as a potential candidate regulating the ECT-mediated glioblastoma malignancy, we investigated the function of PLAGL1 in our glioblastoma edge-TIC models to understand if its targeting holds any translational significance. To this end, we used the two tumor edge-derived glioma sphere models (1051E and 101027E) for lentivirus-mediated gene overexpression (PLAGL1-OE) and knockdown by shRNA (sh#1 and #2). As the control, we used the non-targeting lentiviral construct (Ctrl). Western blotting confirmed both induced overexpression and gene silencing in cells harboring the shRNA construct, with more efficient targeting of PLAGL1 by sh#2 than sh#1 **(Fig. 3A, Supplementary 5.A)**. In both models, PLAGL1-OE displayed significantly higher *in vitro* growth rates, while their growth was largely attenuated by gene silencing of PLAGL1 **(Fig. 3B)**. Using clonal sphere formation as a surrogate *in vitro* indicator of tumor initiating capacity, we found that PLAGL1-OE glioma spheres relatively increased, whereas its gene silencing reduced it with a greater inhibitory effect of sh#2 compared to sh#1 **(Fig. 3C, D, Supplementary Fig. 5B)**. *In vivo* injection of PLAGL1-OE glioma spheres into brains of SCID mice resulted in higher luminescent intensity indicative of their larger tumor sizes by edge-TIC-derived tumor establishment, whereas the shRNA-carrying xenografts displayed significantly lower signals in both of these two glioblastoma edge sphere-derived tumor models **(Fig. 3E)**. Mice with PLAGL1-OE glioma sphere-derived tumors exhibited significantly worse survival with higher tumor burden, while their gene silencing groups displayed improved overall survival with lower tumor burden compared to the control group **(Fig. 3F, Supplementary Fig.6)**. As expected, immunoreactivity to CD109 was strongly correlated with the expression of PLAGL1 in both models **(Fig. 3G)**. Collectively, this data suggested that PLAGL1 regulates the *in vitro* clonality and *in vivo* tumor development originally derived from edge-TICs.

**Figure 3.**
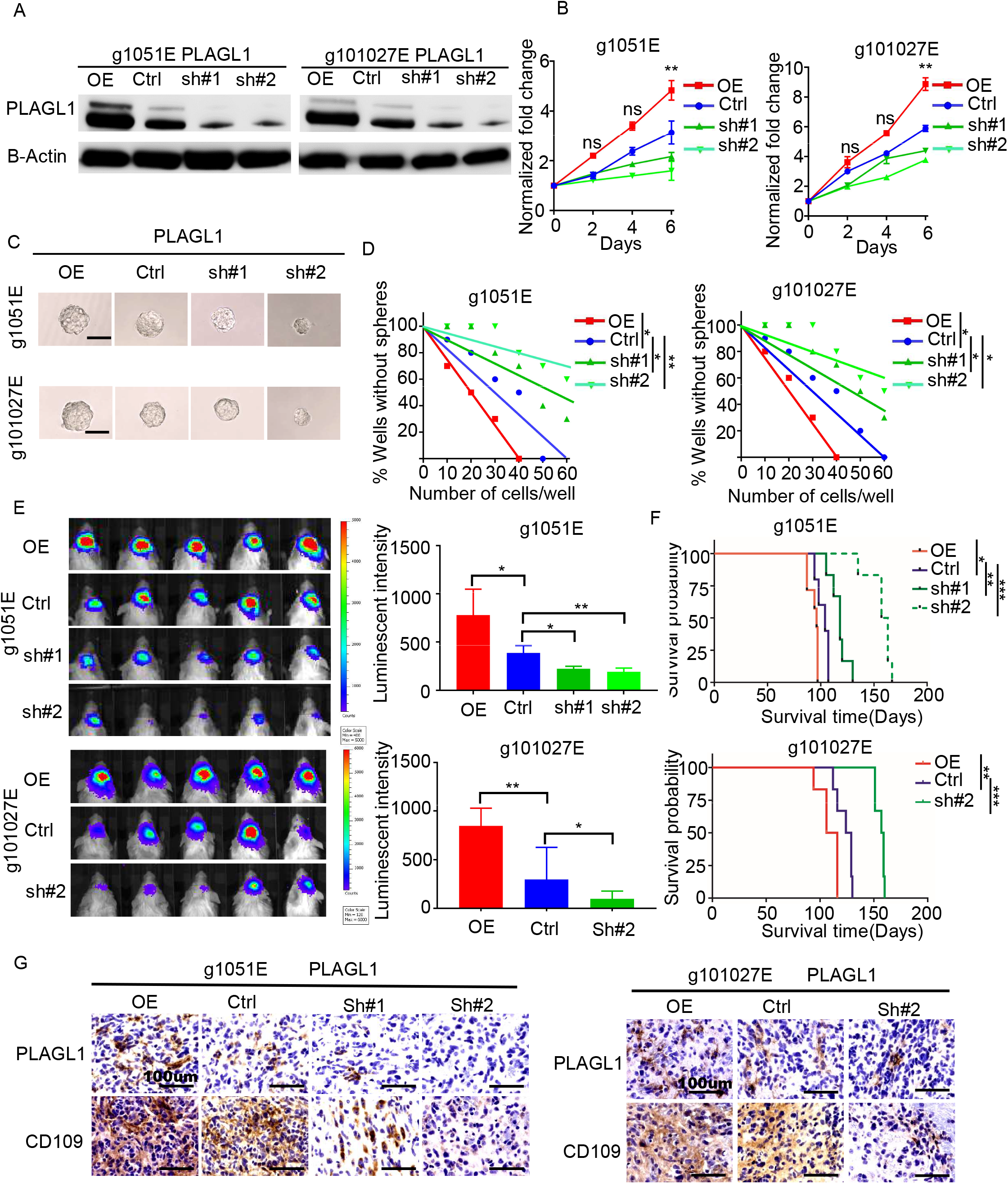
PLAGL1 overexpression enhances, while its silencing diminished, glioblastoma growth in vivo, leading to affect subsequent mouse survival in the edge-TIC models. (A) Western blotting of two patients' tumor edge-derived glioma sphere lines (g1051E and g101027E) after transducing with overexpression vector (OE) or shRNA targeting *PLAGL1* (sh#1 or sh#2) or a nontargeting control (Ctrl). (B) Line charts of *in vitro* growth of the indicated groups (\*\**p*<0.01, n=6, one-way ANOVA). (C) Representative images of the indicated glioma sphere lines after genetic transduction. Scale bar 60 μm. (D) Inverse linear graphs of in vitro clonogenicity assays (limiting dilution neurosphere formation assays) depicting the relationship between PLAGL1 expression and edge-derived GBM spheres (g1051E, g101027E). (\**p*<0.05, \*\**p*<0.01, ELDA analyses) (E) Bioluminescent images (Left) and their quantifications (Right) of orthotopic mouse xenografts established by injection of indicated g1051E and g101027E glioma sphere models. (\**p*<0.05 and \*\**p*<0.01, n=5, one-way ANOVA). (F) Kaplan-Meier analysis of SCID mice harboring intracranial tumors derived from g1051E or 101027E spheres transduced with either overexpressed PLAGL1 (n=7), Ctrl(n=5), shPLAGL1#1(n=6) or shPLAGL1#2 (n=6). \**p</italic><0.05, \*\**p*<0.01, and *** *p*<0.001.* (G) IHC of indicated tumors in SCID mice for CD109 and PLAGL1. Scale bar 100um.

### PLAGL1 binds to the promoter region for CD109 to regulate its transcriptional activity

Lastly, we sought to determine the molecular mechanisms linking the expression of PLAGL1 and CD109. Specifically, we tested if PLAGL1 binds to the promoter region for the CD109 gene in glioblastoma edge-derived cells. Using g1051E spheres, we performed chromatin-immunoprecipitation with the PLAGL1 antibody, followed by qRT-PCR for the CD109 genetic regulatory element that we previously identified as its active promoter^17^ and detected a band indicative of the direct transcriptional regulation of CD109 by PLAGL1 in glioblastoma edge-derived cells. This result was also validated with g101027E cells **(Fig.4A)**. As expected, overexpression of PLAGL1 elevates, while its silencing decreases, the expression of CD109 protein, determined by western blotting in both sphere models **(Fig.4B)**. Collectively, the tightly associated co-expression of PLAGL1-CD109 was, at least in part, mediated through the direct transcriptional regulation of CD109 *via* the TF, PLAGL1.

**Figure 4.**
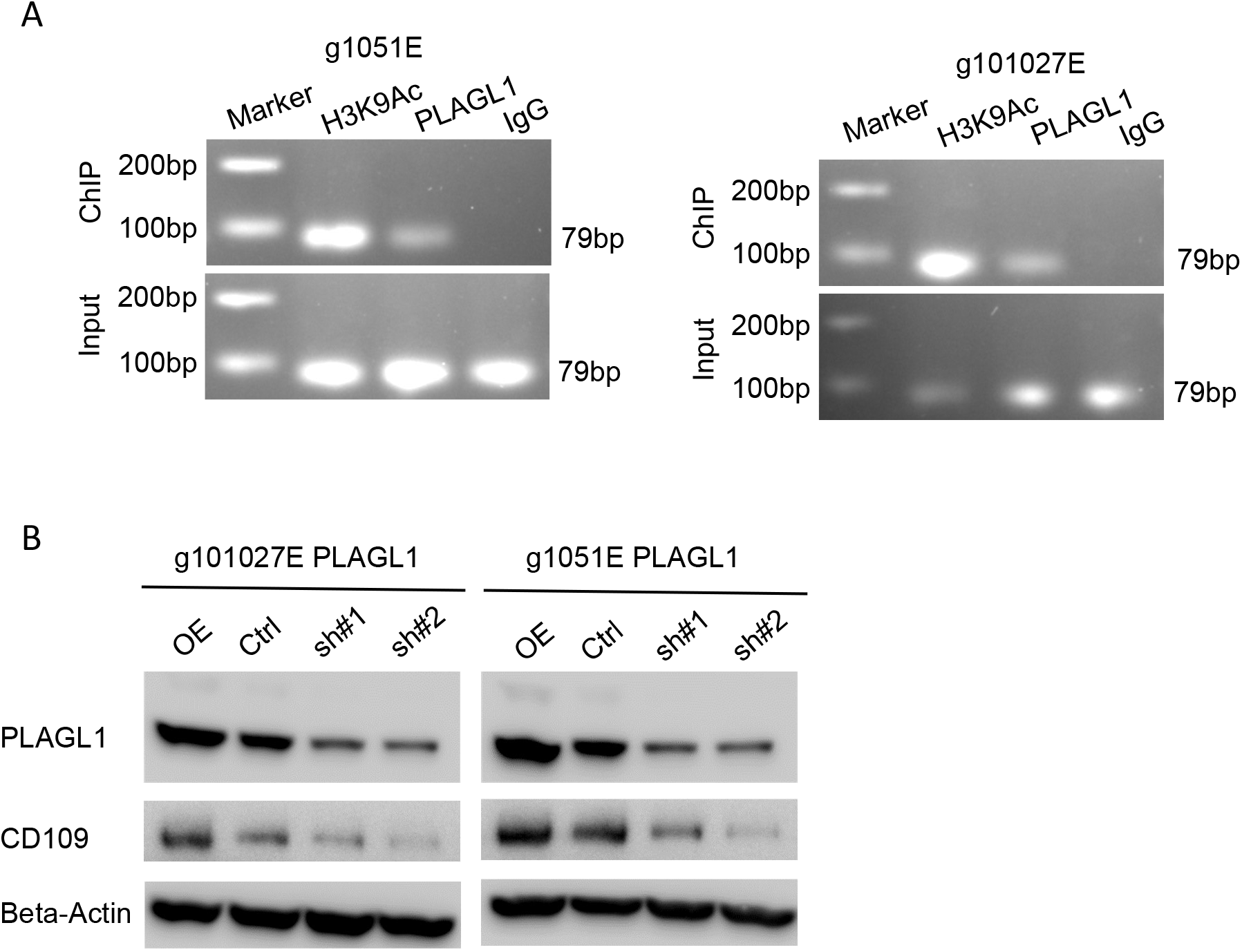
PLAGL1 binds to the promoter region for CD109 to regulate its transcriptional activity. (A) ChIP-qPCR assay showing PLAGL1 binding to the promoter region for the CD109 gene in g1051E and g101027E spheres. H3K9Ac is used as a positive control. (B) Western blotting of CD109 and PLAGL1 in g1051E and g101027E spheres after transducing with overexpression vector, shRNA targeting PLAGL1 (sh#1 or sh#2), or a non-targeting control (Ctrl).

## Discussion

Patients with glioblastoma gain only limited benefit from craniotomy due to the inability to completely eliminate tumor cells from the brain^28,29^. The lethal seeds for tumor recurrence (recurrence-initiating cells) reside predominantly, yet not entirely, at the tumor edge surrounding the resection cavity. In this study, we used CD133 and CD109 expression changes as a reference to indicate ECT. The rationale for this investigation included our previous finding that CD133 and CD109 are preferentially expressed within the TIC subpopulations, selectively within glioblastoma edge- and core-tissues, respectively^17^. While expression of CD109 and CD133 within individual cells in tumors appear to be mutually-exclusive, previous studies indicate that expression of these markers represents a dynamic molecular state^6^. One means of affecting ECT is through radiation, which induces the conversion of edge-associated CD133(+)/CD109(−) cells to the core-associated CD133(−)/CD109(+) cells, thereby developing therapy-resistant tumors *in vivo*. On the other hand, core CD133(−)/CD109(+) cells themselves respond to radiation by secreting factors that promote the radiation resistance of edge-located CD133+)/CD109(−) cells *in vitro* and *in vivo*. Collectively, these prior data suggest the significance of targeting both core- and edge-TICs (marked by CD133 and CD109, respectively) to achieve better outcomes of glioblastoma treatment. However, recent advances in surgical technique, including the imaging-assisted fluorescence-guided surgery in the awake setting, has allowed us to increase the proportion of surgical cases of total or near-total resection of the tumor core lesions. Yet, edge-located tumor cells undoubtedly remain as a key therapeutic target as they are the presumptive sources of recurrent tumors.

In the current study, we examined 37 paired primary-recurrent tumor samples to focus on ECT, validating its association with poorer patient prognoses. It is important to note that both tumor edge and core are composed of tumor cells in all three transcriptomic subtypes, albeit the ratios are slightly different (**Fig. 5**)^18^. Our findings suggest, yet do not definitely prove, relatively weaker correlation of mesenchymal-ness, in comparison to the ECT signature, to patients' poorer prognosis, at least in this patient cohort. This interpretation needs further validation with more clinical evidence, ideally with prospective measurement, from multiple independent groups. The ECT axis could be more clinically-relevant but it remains poorly understood how similar to, or different from, the transcriptomic proneural (and classical)-mesenchymal axis it is. In addition, in many craniotomies, small residual core lesions are left behind. Most likely, they also contribute both directly and indirectly to the tumors to recur, as our recent study suggested^18^. Therefore, we need to be cautious in stating that ECT does not explain all the clinical courses of the primary-to-recurrent glioblastoma progression. More extensive molecular characterization with additional longitudinal case cohorts is warranted.

**Figure 5.**
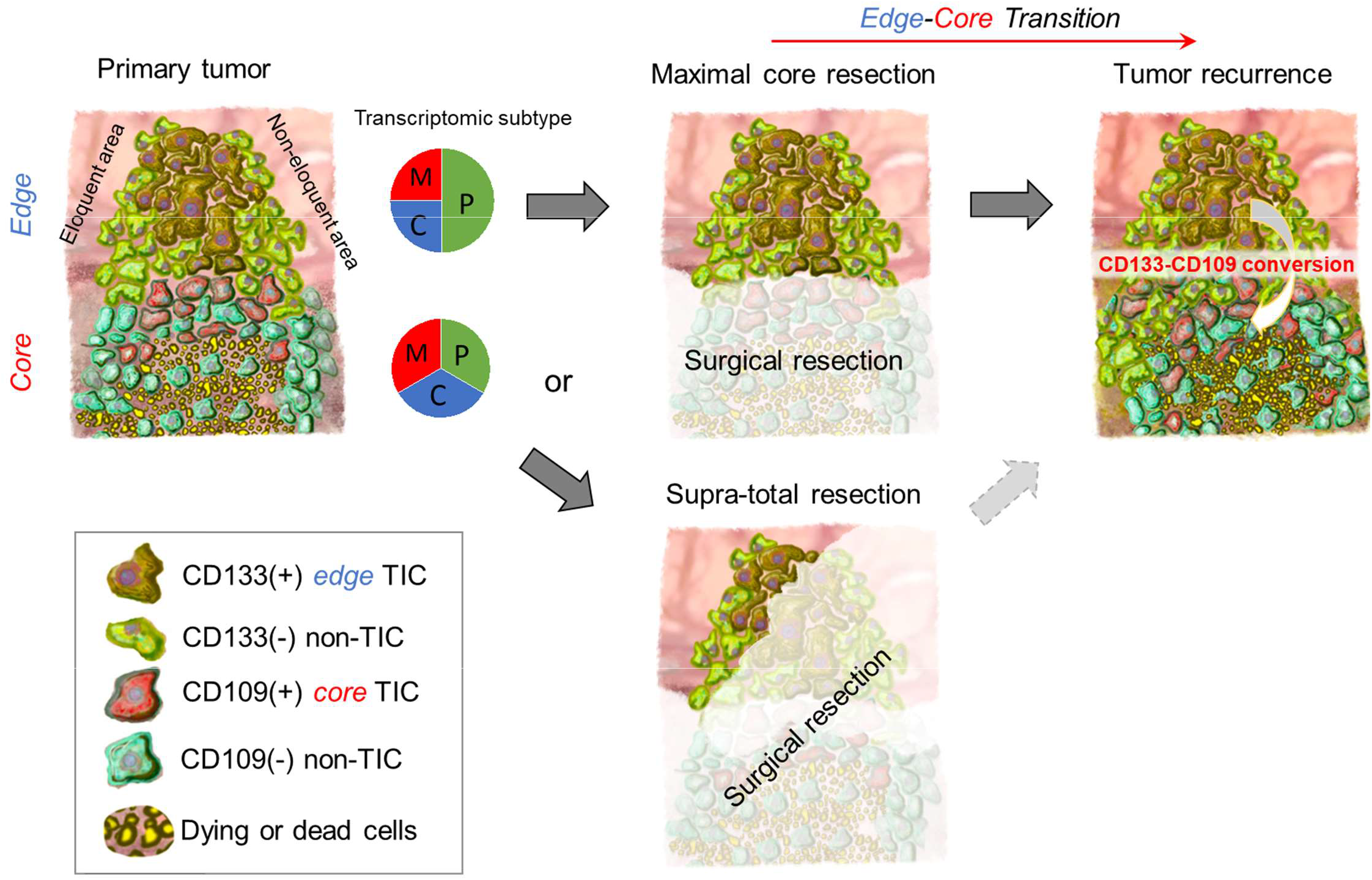
Schematic delineating the edge- and core-located tumor cells in glioblastoma together with intra-tumoral CD133 and CD109 expressions in the TIC subpopulations.

For the phenotypic characterization associated with tumor edge and ECT in glioblastoma, we believe that the recently-established tumor edge- and core-derived glioma spheres represent valuable models, as their xenografts faithfully recapitulate their spatially-distinct tumor lesions in mouse brains. They can allow for the study the tumor recurrence formation from the mixed populations of the core and edge cells^7^. Needless to say, the accuracy of the resected tissue locations within the brains is critical for these spatially-identified models and a number of potential hurdles have to be overcome in order to ascertain these samples (e.g. brain shift, patient safety). In particular, glioblastomas tend to infiltrate into the deep white matter, where a number of functional neuronal fibers run throughout the brain. Obtaining tumor edge tissues from these regions without harming the patients is a critical step in allowing us to establish models that faithfully recapitulate their spatially-distinct pathobiology. Further characterization of our models and developing other tumor edge-reflective models would help facilitate the molecular and phenotypic analyses to identify therapeutic targets in the post-surgical residual tumor cells at tumor edge that subsequently cause patient lethality.

Our data indicated the significance of targeting PLAGL1 to attenuate, yet not completely eliminate, tumor initiation and propagation, accompanied by an impact on survival of tumor-bearing mice. As for its molecular mechanism, we found that this TF directly regulates the ECT gene CD109. Our previous studies demonstrate that CD109 drives ECT, and, thus, by inference, PAGL1 would be expected to do the same. Nonetheless, the role of PLAGL1 in cancer has been controversial. Prior studies have shown that PLAGL1 is a tumor suppressor gene encoding an inducer of apoptosis and cell cycle arrest in various cancers^30,32^ (e.g. breast cancer, hepatoma, colon cancer). Even in glioma, one study has demonstrated the frequent loss of PLAGL1 in their clinical samples. However, another study paradoxically demonstrated a pro-tumorigenic function of PLAGL1 driven by SOX11^33,34^. Here, using pre-clinical models, we provide strong evidence to support the tumorigenic function of PLAGL1 in glioblastoma TICs. In addition, the analysis of clinical samples using public and our own databases were consistent with our experimental findings. PLAGL1-mediated signaling might be context-dependent among various cancer cells, or even within gliomas. Such context-dependency is known to occur in a variety of settings, including ones directly related to these studies. We previously found that histone deacetylase 1 (HDAC1) is a positive transcriptional regulator that drives CD109 gene expression *via* a protein complex formation with an oncogenic TF C/EBPβ, even though HDAC1 is recognized to modulate the compact chromatin structure leading to widespread repression of transcriptional activities in cancers and developmental somatic cells^35,37^. It remains unknown if PLAGL1 forms a larger protein complex with HDAC1 and C/EBPβ in glioblastoma and other cancers.

In conclusion, this study provides a set of clinical and experimental data suggesting the significance of targeting tumor edge-located TICs that subsequently escape current therapies to develop lethal core lesions during tumor recurrence. The PLAGL1-CD109 signaling axis is likely among key drivers for ECT. As the molecular and cellular complexity of glioblastoma is increasingly recognized as a challenging road block to prolong survival of patients, successful removal of tumor core is certainly the mandated first-step, yet it still requires us to learn how to manage the tumor edge in better ways. Further phenotypic characterization of edge-TICs is among key tasks ahead of us to develop effective therapies for glioblastoma.

## Acknowledgments

We would like to express our sincere appreciation to all the patients and families, who kindly allowed us to obtain their tumor samples for this study. We would also thank all our collaborating scientists, as well as the assigned reviewers and Editor for this manuscript, for the constructive comments and suggestions. Lastly, we acknowledge the contribution by all the members in the Nakano and Nam laboratories (past and present) for technical help, including Drs.Seungwon Choi and Harim Koo for MRI acquisition and tumor location annotation works in the Nam laboratory.

## Notes

**Funding** This study was supported by National Institutes of Health (NIH) grants to I.N. (R01NS083767, R01NS087913, R01CA183991, and R01CA201402), a grant from the Korea Health Technology R&D Project through the Korea Health Industry Development Institute (KHIDI), funded by the Ministry of Health & Welfare, Republic of Korea (HI14C3418) to D.H.N., and a grant from the Dr. Miriam and Sheldon G. Adelson Medical Research Foundation to H.I.K.

**Conflict of Interest** The authors report no conflict of interest concerning the materials or the methods used in this study or the findings specified in this paper.

### Competing Interest Statement

The authors have declared no competing interest.

