## Supplementary Figure for "Tumor Edge-to-Core Transition Promotes Malignancy in Primary-to-Recurrent Glioblastoma Progression in a PLAGL1/CD109-mediated mechanism"

### Slide 1
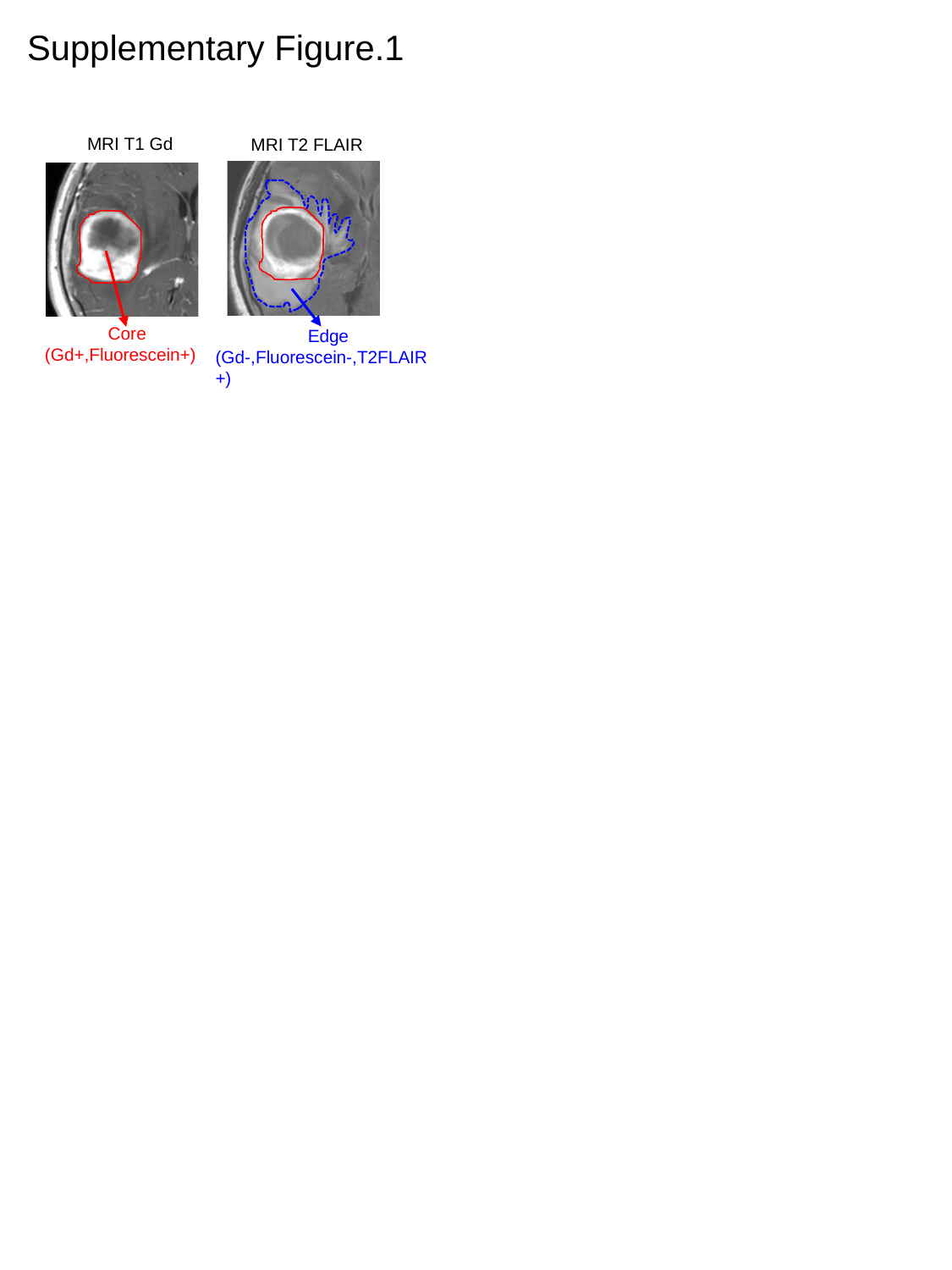

Supplementary Figure.1
 MRI T1 Gd
 MRI T2 FLAIR
 Core
(Gd+,Fluorescein+)
 Edge
(Gd-,Fluorescein-,T2FLAIR +)

### Slide 2
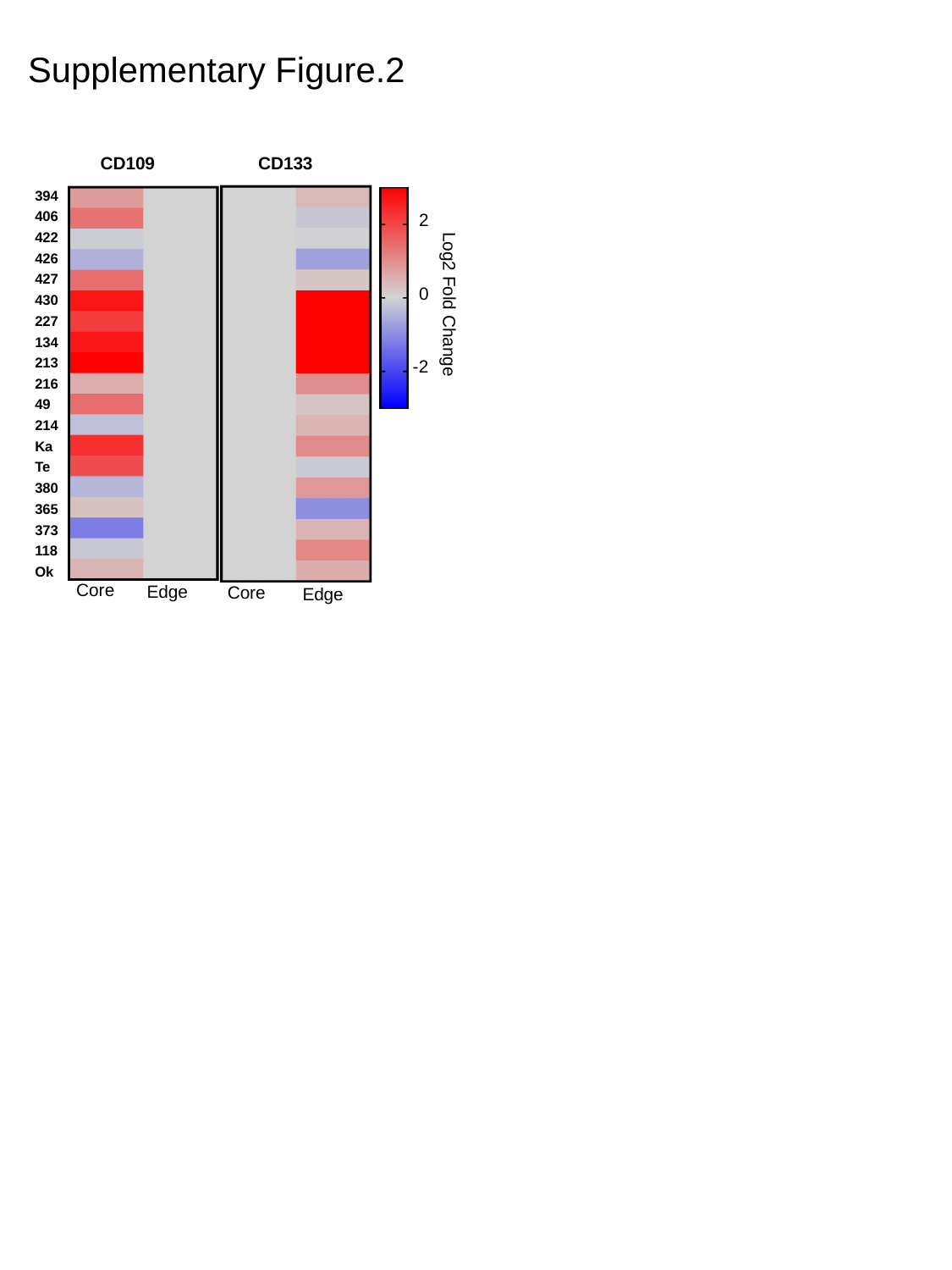

Supplementary Figure.2
CD109
Core
Edge
CD133
Core
Edge
2
Log2 Fold Change
0
-2
| 394 |
| --- |
| 406 |
| 422 |
| 426 |
| 427 |
| 430 |
| 227 |
| 134 |
| 213 |
| 216 |
| 49 |
| 214 |
| Ka |
| Te |
| 380 |
| 365 |
| 373 |
| 118 |
| Ok |

### Slide 3
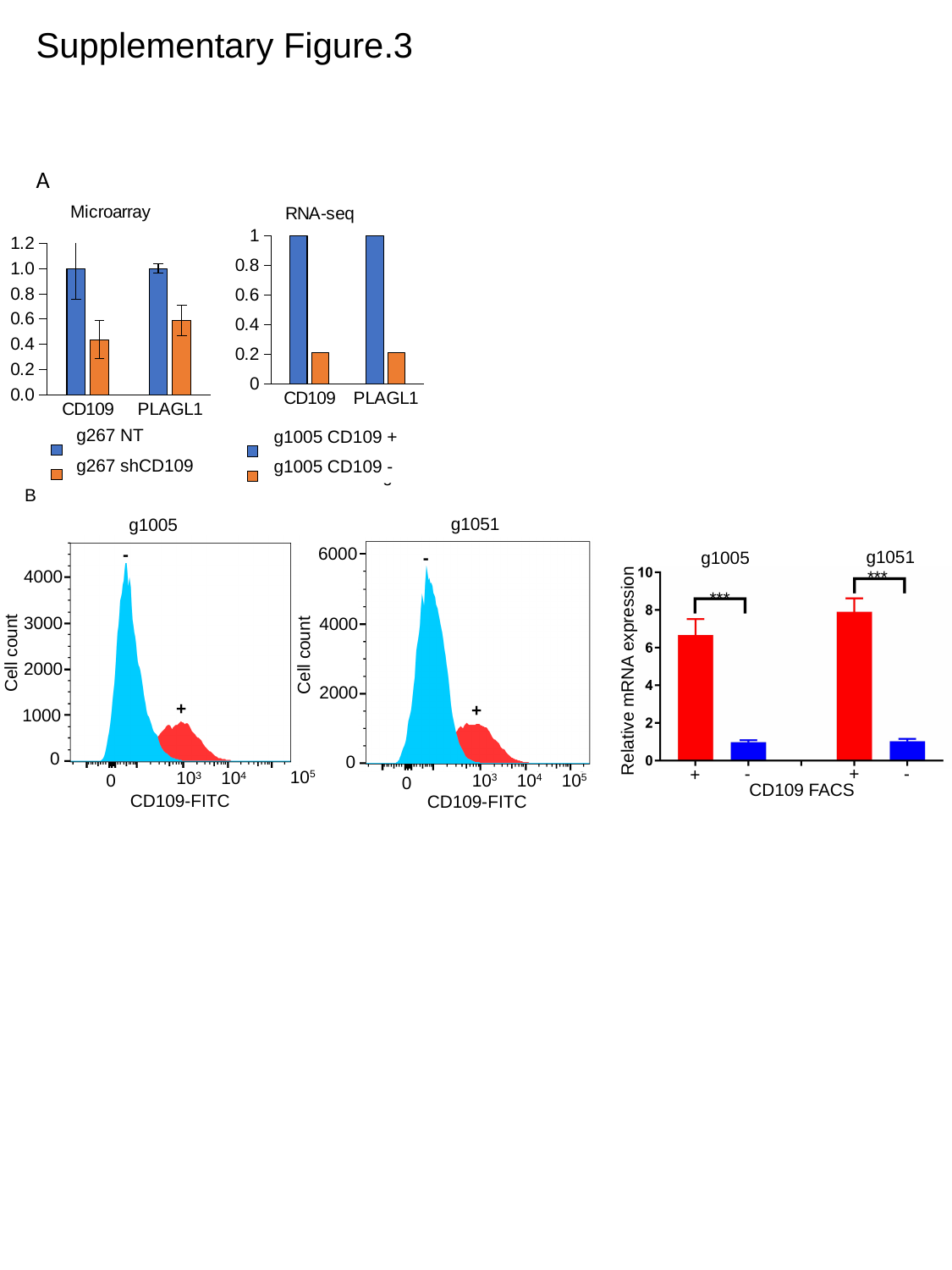

Supplementary Figure.3
A
#### Chart: Microarray
| Category | 267 NT | 267 shCD109 |
|---|---|---|
| CD109 | 1.0 | 0.436313149408814 |
| PLAGL1 | 1.0 | 0.5904348968598082 |
#### Chart: RNA-seq
| Category | 1005 CD109 Pos | 1005 CD109 Neg |
|---|---|---|
| CD109 | 1.0 | 0.21201209851649142 |
| PLAGL1 | 1.0 | 0.20759493670886073 |g267 NT
g1005 CD109 +
g267 shCD109
g1005 CD109 -
B
g1051
6000
4000
2000
0
10^3^
10^4^
10^5^
0
CD109-FITC
Cell count
-
+
g1005
4000
3000
Cell count
2000
1000
0
10^3^
10^4^
0
CD109-FITC
-
+
g1051
g1005
***
***
Relative mRNA expression
-
+
-
+
CD109 FACS
10^5^

### Slide 4
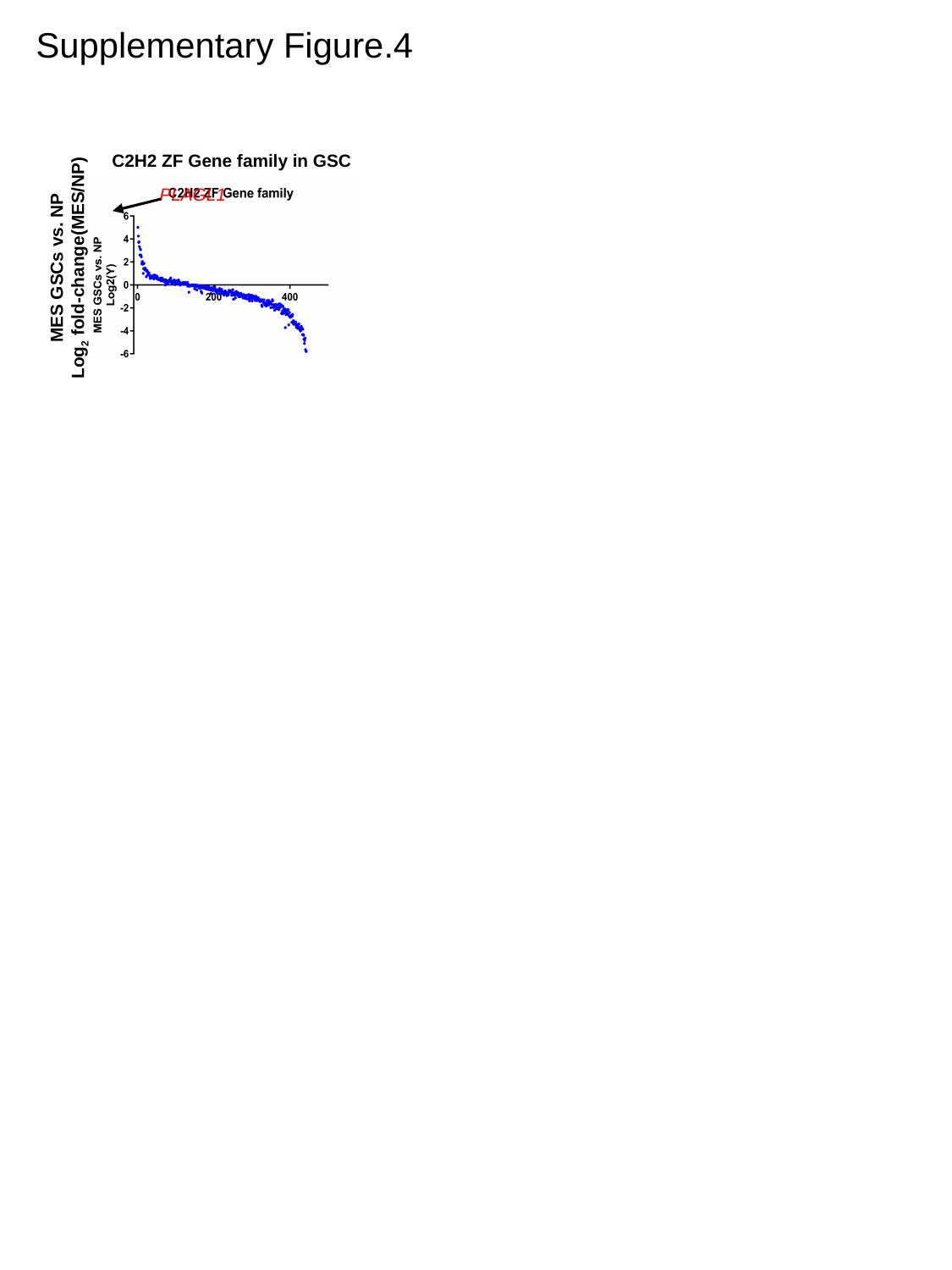

Supplementary Figure.4
C2H2 ZF Gene family in GSC
PLAGL1
MES GSCs vs. NP
Log_2_ fold-change(MES/NP)

### Slide 5
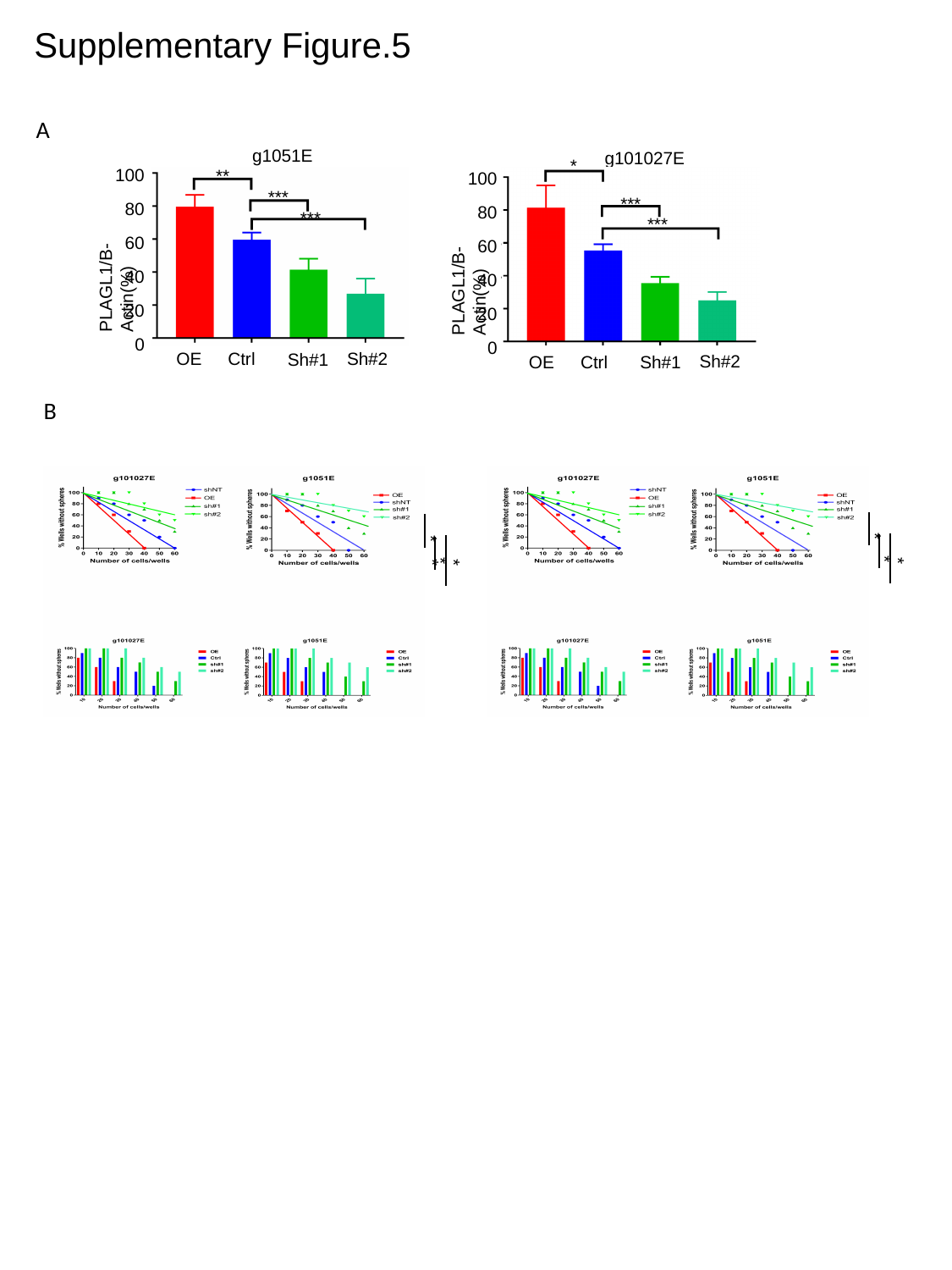

Supplementary Figure.5
A
g1051E
100
80
60
PLAGL1/B-Actin(%)
40
20
0
Sh#2
Ctrl
OE
Sh#1
**
***
***
g101027E
100
80
60
PLAGL1/B-Actin(%)
40
20
0
Sh#2
Ctrl
OE
Sh#1
*
***
***
B
*
*
**
*
*
*

### Slide 6
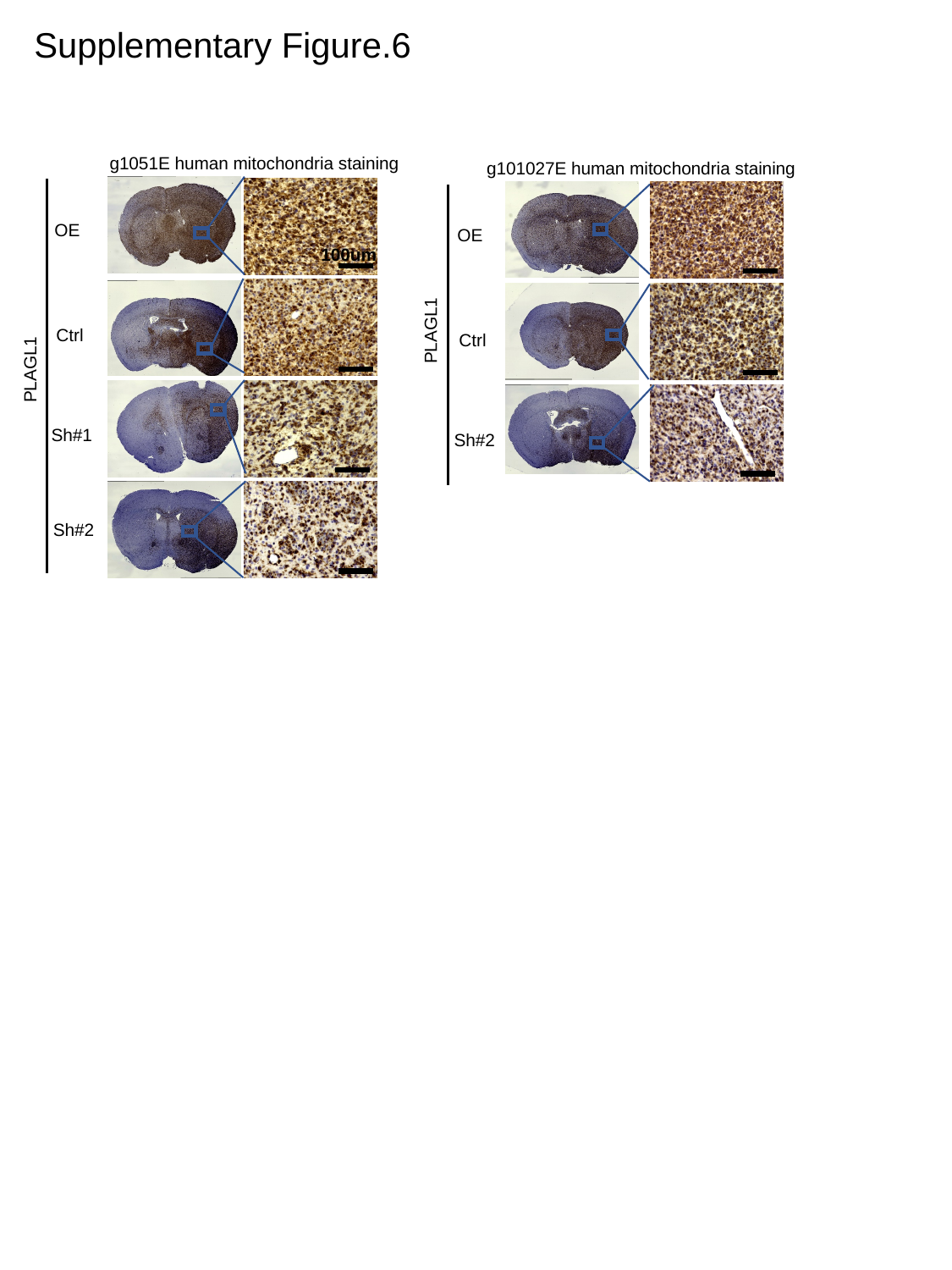

Supplementary Figure.6
g1051E human mitochondria staining
100um
OE
Ctrl
PLAGL1
Sh#1
Sh#2
g101027E human mitochondria staining
PLAGL1
OE
Ctrl
Sh#2

### Slide 7
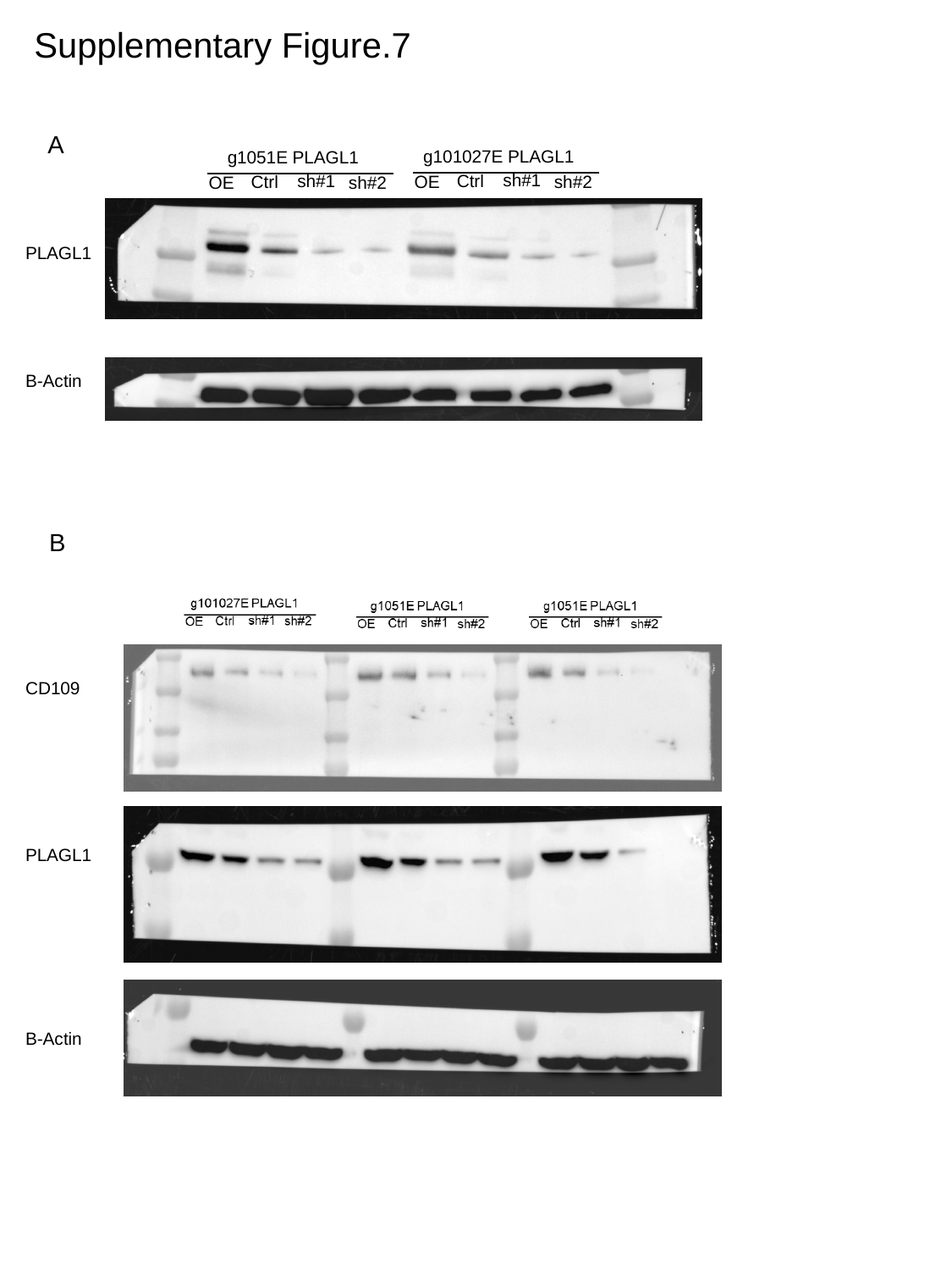

Supplementary Figure.7
A
g101027E PLAGL1
sh#1
Ctrl
sh#2
OE
g1051E PLAGL1
sh#1
Ctrl
sh#2
OE
PLAGL1
B-Actin
B
CD109
PLAGL1
B-Actin
