## Supplementary material for "Tumor Edge-to-Core Transition Promotes Malignancy in Primary-to-Recurrent Glioblastoma Progression in a PLAGL1/CD109-mediated mechanism": Legends of supplementary Figure

**Supplementary Figure 1**

Representative MRI images (T1-weighted image with Gadolinium enhancement (left) and T2/FLAIR image (right)) to define the glioblastoma core and edge lesions. Fluorescein is used during surgery to indicate the tumor tissues in the non-enhancing FLAIR abnormal edge lesions.

**Supplementary Figure 2**

mRNA expression profile of *CD133(Prominin)* and *CD109* in paired glioblastoma edge and core samples (n=19, *p*=0.005 for *CD109*, *p*=0.018 for *CD133*, paired *t*-test.)

**Supplementary Figure 3**

**(A)** *CD109* and *PLAGL1* expressions in g267 glioma spheres transduced with shNT and shCD109 determined by cDNA Microarray (n=3) (left). *CD109* and *PLAGL1* expressions in CD109-positive and -negative cells in g1005 glioma spheres sorted by FACS determined by RNA-sequencing (seq) (n=1).

**(B)** Cell sorting with g1005 and g1051 glioma spheres for separation of CD109-positive and -negative cells (left) and subsequent qRT-PCR confirmation for the expression of *CD109* in the FACS sorted cells (g1053 and g0573). Data are means ± SD (n=3). ****p*<0.01.

**Supplementary Figure 4**

mRNA expression levels of the C2H2 TF family genes in MES glioma sphere lines compared to the neural progenitor cells(NP) indicates *PLAGL1* is the second highest expressor in MES cells.

**Supplementary Figure 5**

**(A)** Graph indicating the quantification of the Western blot images shown in Fig. 3A. (**p*<0.05, ***p*<0.01, and *** *p*<0.001)

**(B)** Bar graphs of the limiting dilution glioma sphere forming assay, depicting the relationship between PLAGL1 expression and clonal populations in the edge-derived glioma spheres (g1051E, g101027E). (**p*<0.05, ***p*<0.01, ELDA analyses)

**Supplementary Figure 6**

IHC for human-specific mitochondria with mouse brains harboring intracranial tumors derived from g1051E spheres transduced with overexpressed PLAGL1, Ctrl, shPLAGL1#1, or shPLAGL1#2, and from g101027E spheres transduced with overexpressed PLAGL1, Ctrl, or shPLAGL1#2. Higher magnifications (right) exhibit the difference in cellular densities between the compared groups. Scale bar 100um.

**Supplementary Figure 7**

(A) Raw images for WB in Figure 3A.
(B) Raw images for WB in Figure 4B.
