## Supplementary material for "Tumor Edge-to-Core Transition Promotes Malignancy in Primary-to-Recurrent Glioblastoma Progression in a PLAGL1/CD109-mediated mechanism": Methods supplement

**Supplementary Methods**

Most of the methods described below are described in our previous papers in detail^1-7^. All the experiments were performed as at least three independent procedures, unless described otherwise.

*Data reliability and validity*

All the experiments were performed by two or more researchers involved in each procedure. The in vitro experiments were repeated at least three independent procedures to obtain valid data. The clinical data in Table 1 and Figure 1 were collected in the Nam laboratory and the data was discussed between C.L., H. J. C., D. H. N., and I.N. The data in Fig. 2-4 were primarily gathered in the Nakano laboratory and their interpretation was discussed and confirmed among H.I. K., D.H. N, and I.N. The manuscript was primarily written by I.N. with extensive discussion with H.I.K. and D.N. All the raw data are placed in Supplementary Figure 8. After completion of the entire manuscript and prior to submission to the journal, the corresponding authors (D.H. N. and I.N.) confirmed the agreement of everything with all the involved authors in this study.

*Delta value*

Delta value was defined as:

Delta-X=log2(($\frac{Recurrent RPKM+1 of X gene}{Primary RPKM+1 of X gene})$, Which reflects the fold changes of gene expressions after recurrence.

*Glioblastoma Transcriptomic Subtyping*

To estimate original glioblastoma subtype and tumor-intrinsic glioblastoma subtype from C-GBM samples, we evaluated single sample GSEA (ssGSEA) scores for each subtype markers^10,11^. Each subtype score was Z-normalized with in-house 275 glioblastoma reference set for C-GBM samples, and the subtype with highest Z-score within a sample was assigned as its subtype of each sample.

*RNA isolation and Quantitative Real-Time PCR (RT-qPCR)*

Total RNA was extracted using the RNeasy mini kit (QIAGEN) according to the manufacturer’s instructions. RNA concentration was determined using Nanodrop One (Thermo Scientific). cDNAs were synthesized using iScript reverse transcription supermix (Bio-Rad) according to the manufacturer’s protocol. RT-qPCR analysis was performed on StepOnePlus thermal cycler (Thermo scientific) with SYBR Select Master Mix (Thermo scientific). *GAPDH* mRNA was used as an internal control. Primer sequences are shown in Supplementary Table 3.

*Immunohistochemistry (IHC)*

IHC were performed as has been previously described^1-7^. Tumors embedded in paraffin blocks were deparaffinized and hydrated *via* progression through ethanol series. After microwave-mediated antigen retrieval in IHC antigen retrieval solution pH 6 (Invitrogen, Carlsbad, CA), slides were incubated in 0.3% hydrogen peroxide solution in methanol for 15 minutes at room temperature (RT) to inhibit intrinsic peroxidase activity. Next, samples were blocked with serum-free protein block solution (Thermo Fisher Scientific, Rockford, IL) and incubated with the corresponding primary antibodies (PLAGL1 ab181457(abcam)，CD109 SC271085 (Santa Cruz Biotehcnology)), overnight at 4°C. Slides were then stained with Signal Stain Boost IHC Detection Reagent (Cell Signaling Technology, Beverly, MA) and visualized with DAB peroxidase substrate kit (Vector Laboratories, Inc.; Burlingame, CA). Images were captured using an EVOS® FL microscope (Advanced Microscopy Group, Bothell, WA).

*Glioma sphere cultures*

Glioma sphere cultures from clinical samples were cultured in DMEM/F12 medium containing 2% B27 supplement (% vol), 2.5 mg/ml heparin, 20 ng/ml bFGF, and 20 ng/ml EGF, as described previously^1-7^. The bFGF and EGF reagents were added twice a week, and the culture medium was replaced every 7 days. Experiments with these glioma spheres were performed with lines that were cultured for fewer than 40 passages from their initial establishment.

*Western blot analysis*

Cells were lysed for 30 min on ice in RIPA buffer (Sigma) containing 1% protease and 1% phosphatase inhibitor cocktail (Sigma). Lysates were pre-cleaned by centrifugation at 160 000g, 15 min, 4 °C. Protein concentration was determined by Bradford method. Equal amounts of protein lysates (30 μg/lane) were fractionated by NuPAGE Novex 4-12% Bis-Tris Protein gel (Thermo scientific) and transferred to a PVDF membrane (Thermo scientific). Subsequently, the membrane was blocked with 5% Blotting Grade Blocker Nonfat Dry Milk (Bio-Rad) for 1 hour and then incubated with corresponding primary antibody overnight and next incubated with peroxidase conjugated secondary antibodies (GE Healthcare) for 1 hour. Staining was visualized with Amersham ECL Western Blot System (GE Healthcare). The used antibodies are listed in Supplementary Table 4.

In vitro *cell growth assay*

The cell numbers of glioma spheres with lentiviral gene transduction were determined using the Alamar Blue assay (Thermo Fisher Scientific, Rockford, IL), as described previously^1-7^. Briefly, cells were seeded at the density of 3,000 cells per well in 96-well plates (excitation at 515-565nm, emission at 570-610nm were the parameters employed); a Synergy HTX multi-mode reader (BioTek; Winooski, VT) was utilized for quantification in all experiments.

*Neurosphere formation assay*

1051E and 101027E glioma spheres were seeded into 96-well plates at 10, 20, 30, 40, 50, and 60 cells per well. After 10 days for 1051E cells and 12 days for 101027E cells, the numbers of spheres with diameters greater than 60 μm were counted. Data were analyzed as described previously^12^ (http://bioinf.wehi.edu.au /software/elda/;).

*Lentivirus Production and Transduction*

Lentiviral vectors expressing shRNA for PLAGL1 (sh#1 and sh#2) were purchased from Sigma. Plasmid DNA was purified with HiSpeed Plasmid Midi Kit (Qiagen). For lentiviral production, HEK293FT cells were transfected with the vectors (Sigma) and two packaging plasmids psPAX2 and pMGD2) using the CalPhos Mammalian Transfection Kit (Clontech) according to the manufacturer’s protocol. The lentiviral particles were harvested 72 h after transfection and were concentrated 100-fold using a Lenti-X concentrator (Clontech) and stocked -80°C until infection. One day before infection of glioma spheres were dissociated into single cells with accutase and seeded on laminin coated 6-well plates, 5×10^5^ cells per well. Next day these infected cells were incubated with viral supernatants for 24 hours in the presence of 8 µg/ml polybrene (EMD Millipore). At 24 h after infection, medium was replaced and cells were collected at 5 days after infection.

*Targeting sequences for PLAGL1*

PLAGL1 (#1:TRCN0000021236 Clone ID:NM_002656.2-3144s1c1, sequence: CCGGGCCTCATTTCCATCATGCATTCTCGAGAATGCATGATGGAAATGAGGCTTTTT; #2:TRCN0000021238 Clone ID : NM_002656.2-1981s1c1, sequence: CCGGGAGAAGACGTTCAACCGGAAACTCGAGTTTCCGGTTGAACGTCTTCTCTTTTT). For the overexpression experiment, TetO-FUW-PLAGL1 was a gift from Rudolf Jaenisch^13^ (Addgene plasmid # 61542) and doxycycline was used for the induction.

*Chromatin Immunoprecipitation Assay (ChIP)*

ChIP assay was performed after cross-linking cells using formaldehyde. DNA was sonicated using an Ultrasonic Processor (GE130, Sorvall) at 12 cycles of 30 seconds with a 30 seconds interval between cycles. Sonicated DNA was then centrifuged at 13, 5000 rpm at 4°C. Supernatant from 100,000 cells was used for each ChIP assay using MAGnify ChIP system (Invitrogen). Two micrograms of mouse IgG, PLAGL1 (Abcam-ab 129063) H3k9ac (CST,9649s) was used per ChIP. Immunoprecipitated DNA was analyzed by electrophoresis. Primer sequences are shown in Supplementary Table 2.

*In vivo Intracranial Xenograft Tumor Models*

As described previously^1-7^, SCID mice (6-8 weeks old) were used for intracranial implantation of the used glioma sphere lines. The glioblastoma cell suspensions (5×10^5^ cells in 5 μl of PBS) was injected into brains of SCID mice as previously described^1-7^. When neuropathological symptoms were developed due to tumor burden, mice were euthanized and perfused with ice-cold PBS and 4% paraformaldehyde (PFA). These mouse brains were dissected, fixed in 4% PFA for 24 hours, and then transferred to 10% formalin for IHC.

In vivo *Bioluminescent Imaging*

To monitor tumor growth in living animals, glioma spheres were transduced with lentiviral particles (pHAGE PGK-GFP-IRES-LUC-W) for co-expression of green fluorescence protein (GFP) and luciferase, and then GFP-expressing cells were sorted by FACS. Glioma spheres expressing GFP/luciferase were intracranially transplanted into SCID mice. To examine the tumor growth, animals were administrated intraperitoneally with 2.5 mg/100ul solution of XenoLight D-luciferin (PerkinElmer) and anesthetized with isoflurane for the imaging analysis. The tumor luciferase images were captured by using an IVIS 100 imaging system (PerkinElmer).
