## Supplementary material for "Tumor Edge-to-Core Transition Promotes Malignancy in Primary-to-Recurrent Glioblastoma Progression in a PLAGL1/CD109-mediated mechanism": table1

|  | CD133^down^/CD109^up^ | Others | *p*-value |
| --- | --- | --- | --- |
| No. of patients | 15 | 22 |  |
| Age | 54.0 ± 10.2 | 48.2 ± 9.3 | NS |
| Gender(M/F) | 8/7 | 13/9 | NS |
| % of distant recurrence | 33.3% (5/15) | 27.3% (6/22) | NS |
| Radiotherapy between 1^st^ and 2^nd^ surgery (%) | 100% (15/15) | 95.5% (21/22） | NS |
| Chemotherapy between 1^st^ and 2^nd^ surgery (%) | 80% (12/15) | 95.5% (21/22) | NS |
| Delta-CD133 | -1.19 ± 0.98 | 0.21 ± 0.85 | 5.02E-5 |
| Delta-CD109 | 0.93 ± 0.76 | -0.20 ± 0.77 | 9.58E-5 |

**Table 1.** Demographics and clinical characteristics of the patients in this study.

NS: not significant.
