## supplementary.table.2 for "Tumor Edge-to-Core Transition Promotes Malignancy in Primary-to-Recurrent Glioblastoma Progression in a PLAGL1/CD109-mediated mechanism"

| ADAMTS2 | GZMB | RASSF3 |
| --- | --- | --- |
| ANXA4 | ISLR | RBPMS |
| BNC2 | KIAA1217 | SNX9 |
| CD109 | KIF16B | STX3 |
| CD55 | LURAP1L |  |
| CFB | MYLIP |  |
| COL1A1 | MYO1D |  |
| CPPED1 | PERP |  |
| FOXP1 | PLAGL1 |  |
| GALNT6 | POPDC3 |  |
| GCNT1 | PTPRM |  |

Table. 26 up regulated gene in CD133^down^/CD109^up^ group
