## supplementary.table.3 for "Tumor Edge-to-Core Transition Promotes Malignancy in Primary-to-Recurrent Glioblastoma Progression in a PLAGL1/CD109-mediated mechanism"

| **Name** | **Sequence** |
| --- | --- |
| PLAGL1 FW | AAAGATGCTTCTACACCCGGA |
| PLAGL1 RV | AGTGGGTCTTCTTGGTATGCC |
| CD109 FW | AAGCCAGTGAAAGGAGACGTA |
| CD109 RV | CCAGGGGAAGATAGATCCAGG |
| CD133 FW | AGTCGGAAACTGGCAGATAGC |
| CD133 RV | GGTAGTGTTGTACTGGGCCAAT |
| GAPDH FW | GAAGGTGAAGGTCGGAGTCA |
| GAPDH RV | TTGAGGTCAATGAAGGGGTC |
| CD109 ChIP FW | CAGTGCGAGTTCTCTTCTTCTT |
| CD109 ChIP RV | GAGGTCTACTGCTTTCCTTTCC |

Table.3 List of primers used in this study.
