## supplementary.table.4 for "Tumor Edge-to-Core Transition Promotes Malignancy in Primary-to-Recurrent Glioblastoma Progression in a PLAGL1/CD109-mediated mechanism"

Table.4 Antibodies used in this study.

| **Antibodies** | **Source** | **Cat.No** | **Application** |
| --- | --- | --- | --- |
| PLAGL1 | Abcam | ab129063 | WB，ChIP |
| PLAGL1 | Abcam | ab181457 | IHC |
| H3k9ac | Active motif | 39137 | ChIP |
| CD109 | Santa Cruz  Biotechnology | sc-271085 | WB,IHC |
| β-ACTIN | Cell Signaling Technology | 3700S | WB |
| Anti-Rabbit 2nd Ab | Millopore | GENA934 | WB |
| Anti-Mouse 2nd Ab | Millopore | GENA931 | WB |
